# *In vivo* pulse-labeling of isochronic cohorts of cells in the central nervous system using FlashTag

**DOI:** 10.1101/286831

**Authors:** Subashika Govindan, Polina Oberst, Denis Jabaudon

**Affiliations:** Department of Basic Neurosciences, University of Geneva, Geneva, Switzerland; Clinic of Neurology, Geneva University Hospital, Geneva, Switzerland; Interfaculty Institute of Bioengineering, EPFL, Lausanne, Switzerland

## Abstract

This protocol describes a fluorescence birthdating technique to label, track and isolate isochronic cohorts of newborn cells in the central nervous system *in vivo*. Injection of carboxyfluorescein esters into the cerebral ventricle allows pulse-labeling of M-phase progenitors in touch with the ventricle and their progeny across the central nervous system, a procedure we termed FlashTag. Labeled cells can be imaged *ex vivo* or in fixed tissue, targeted for electrophysiological experiments, or isolated using Fluorescence-Activated Cell Sorting (FACS) for cell culture or (single-cell) RNA-sequencing. The dye is retained for several weeks, allowing labeled cells to be identified postnatally. This protocol describes the labeling procedure using *in utero* injection, the isolation of live cells using FACS, as well as the processing of labeled tissue using immunohistochemistry.

## INTRODUCTION

The ability to track neurons from their time of birth until adulthood is crucial to resolve neuronal lineages and investigate the developmental mechanisms underlying neuronal diversity. Neurons in the central nervous system (CNS) are born within two germinal zones, a ventricular zone (VZ), which is located adjacent to the ventricular wall and contains radial glia, and, in many brain regions, a subventricular zone (SVZ), which is located more basally and contains intermediate progenitors^1^. Within the VZ, and across the CNS, radial glial cells undergo interkinetic migration^1^, such that their soma is at a different location at different phases of the cell cycle: S-phase occurs at the basal-most border of the VZ, while M-phase (cell division) occurs at the ventricular wall, in contact with the ventricular cavity, which is thus where neurons are born (Fig 1a). In the present protocol, we describe a procedure in which we take advantage of the juxtaventricular location of M-phase progenitors to pulse-label them by intraventricular injection of a carboxyfluorescein ester (a procedure we termed “FlashTag”, FT). This procedure allows fluorescent tagging of VZ progenitors and tracking of time-locked cohorts of their postmitotic progeny, including neurons, throughout corticogenesis and early postnatal development.

**Fig 1.**
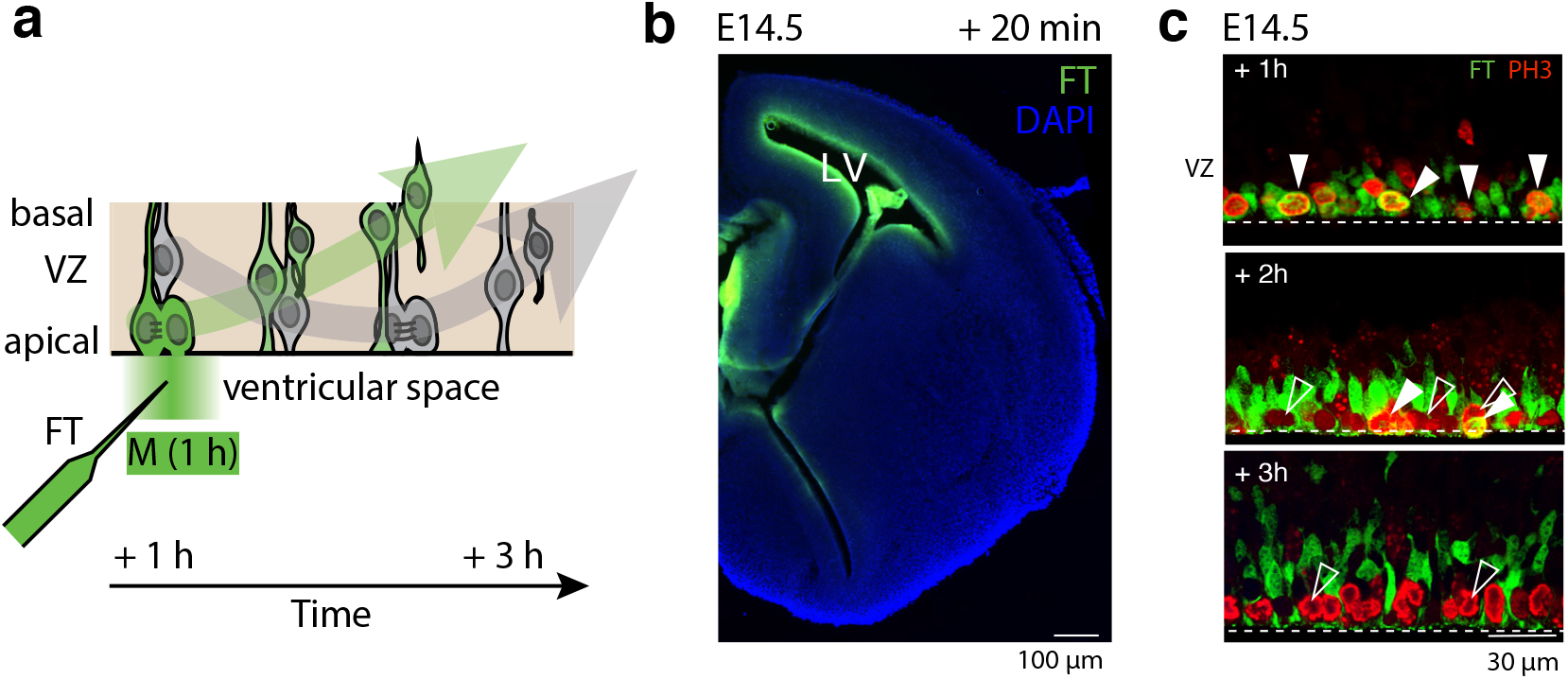
FT specifically labels M-phase VZ progenitors. **(a)** FT specifically labels ventricular zone (VZ) progenitors as they undergo mitosis (M) in contact with the ventricular border, resulting in labeling of daughter cells. **(b)** Twenty minutes after labeling strong fluorescence is observed along the entire ventricular wall. LV: Lateral ventricle. **(c)** FT labels mitotic cells at the ventricular border, as identified by the expression of the mitosis-specific marker PH3 in labeled cells. Labeling is restricted to a period of 1 – 2 hours, and cells born 3 h after labeling (PH3^+^) are FT^-^. Illustrations from ref. 7.

### Birthdating techniques: an overview

The first-developed and probably still most widely used technique to permanently label neurons based on their embryonic time of birth is 5-bromo-3′-deoxy-uridine (BrdU) birthdating^2,3^. BrdU is a thymidine analogue that is incorporated into cells during DNA synthesis, *i.e.* during S phase. Administration of BrdU (usually through intraperitoneal (i.p.) injection in pregnant dams or *via* drinking water) results in systemic labeling of all S-phase cells in the embryo, with no spatial restriction, such that in the CNS, both VZ and SVZ-born neurons are labeled. Of note, the term “birthdating” is somewhat of a misnomer, because BrdU-labeled daughter neurons are not neurons which were born at the time of injection (which would be the case upon M-phase labeling, as occurs with FT), but instead are the progeny of cells which were undergoing S-phase at the time of labeling. Following i.p. injection in pregnant mice, BrdU labeling is detected within 30 minutes, and dividing cells incorporate BrdU during a time window of approximately 2-6 hours^2,4^. In order to reveal BrdU labeling, tissue fixation and an aggressive DNA denaturation protocol is required^3^, such that BrdU birthdating cannot be combined with applications in which live cells need to be isolated or analysed, including live imaging, electrophysiology or transcriptomics. Other thymidine analogues, such as 5-ethynyl-2-deoxyuridine (EdU), 5-chloro-2’-deoxyuridine (CldU) and 5-iodo-2’-deoxyuridine (IdU) exist, which can be used for combinatorial labeling of cells born at different times, but which suffer from the same limitations^5^. Finally, it is important to note that BrdU is also incorporated during nucleotide replacement during DNA repair^6^ (which occurs during early neuron differentiation^7,8^) such that it is not an entirely specific marker of S phase. Results of BrdU experiments should thus be examined with caution depending on context.

Another second birthdating technique that partially overcomes the limitations of BrdU is viral labeling. This approach relies on the use of a replication-deficient virus (typically a retrovirus) expressing a reporter protein^9^. The virus is injected into the cerebral ventricles of developing embryos, from where it infects cells that are located at the ventricular border. Upon infection, the virus integrates into the genome and initiates expression of a reporter protein. While this approach is applicable to *in* and *ex vivo* applications, such as live imaging, and infection is limited to a specific cohort of cells (even allowing clonal analysis when very low titers of viruses are used, see ref.^10^), a major limitation is that it takes a considerable amount of time until the fluorescent reporter is expressed (usually about a day or more in the mouse^11^), such that the precise time of birth of the labeled neurons is difficult to determine without using additional approaches (such as BrdU birthdating). This limitation makes it difficult to use viral labeling to analyse cells immediately after their birth^9,12^.

A third commonly used approach to label newborn cells in the developing brain is *in utero* electroporation. In this procedure, a plasmid coding for a reporter protein is injected into the ventricle and electrical pulses are applied to enable entry of the plasmid into cells. Plasmid transfer is most efficient in M-phase cells, but also affects S-phase cells and is not restricted to the ventricular border^13,14^ (see in particular Supplementary Fig.1 in ref.14 in which a BrdU-labeled plasmid was used to identify cells exposed to the plasmid immediately after electroporation, revealing that all VZ and SVZ cells are exposed). Thus, a broad range of unsynchronized progenitors is labeled, which makes analysis of the progeny complex, in terms of developmental trajectories and lineage relationships. Another limitation of *in utero* electroporation is that detection of labeled cells is possible only after expression of the reporter protein (typically around 10 hours in the mouse^14^), making it difficult to study events occurring upon or immediately following cell division.

To circumvent the limitations of the birthdating techniques described above, we recently developed FT to birthdate, track and isolate isochronic cohorts of VZ-born cells in the developing central nervous system^7^. This technique takes advantage of the fact that VZ progenitors divide exclusively at the ventricular wall, where they can be specifically labeled, and allows labeling a spatially defined cohort (cells in the VZ across the CNS) of cell cycle phase-locked cells (M-phase) with a high temporal resolution (Fig 1): an intracellular fluorescent signal is detected within 20 minutes. Furthermore, at least in the neocortex, labeling is restricted to 1-2 hours (i.e. approximately 1-2 round of mitosis) which follow injection (Fig 1c). The fluorescence signal can be detected without immunohistochemistry, allowing FT to be used for *ex vivo* (and, in principle, *in vivo*) live imaging and electrophysiology, as well as to isolate progenitors and neurons for cell culture or bulk / single-cell RNA sequencing.

### FT principle and direct comparison with BrdU and in utero electroporation

Here, we will briefly review the use of FT for neuronal birthdating and compare it to other existing strategies (see Table 1 and Box 1). To perform FlashTag, carboxyfluorescein diacetate succinimidyl ester (CFSE) is injected into the ventricular system, resulting in the specific labeling of mitotic cells at the ventricular border, which are in contact with the cerebrospinal fluid (Fig 1a). CFSE is colourless when it is extracellular and becomes fluorescent only after being enzymatically processed inside the cell (see Box 1 for details), such that 20 minutes after injection, high intracellular fluorescence is observed along the entire ventricular walls (Fig 1b). While weak fluorescence can be observed in cells further away from the ventricular border (most probably due to diffusion), only cells directly abutting the ventricles show high levels of fluorescence. Once injected in the ventricle, FT labeling occurs for a time window of 1-2 hours: while 1 h after labeling all mitotic progenitors at the ventricular border (identified by expression of the mitosis-specific marker phospho-histone H3 (PH3)) are labeled with FT, 2 hours after labeling most new mitotic cells are not FT^+^ and 3 hours after injection none of the PH3^+^ cells at the ventricular border are FT^+^, and FT^+^ cells have now moved away from the ventricular border (Fig 1c).

**TABLE 1.** Comparison of FlashTag with BrdU, intraventricular viral labeling and electroporation

| Parameter | BrdU labelling | Intraventricular viral labeling | Electroporation | FlashTag |
| --- | --- | --- | --- | --- |
| Time window of labeling | 2 - 6 hours | Probably a few hours | Immediate | < 3 hours |
| Time after which labeling can be detected | Approx. 30 min | < 24 h | Approx. 10 h | Approx. 20 min |
| Cell cycle phase of labeling | S-phase | M-phase | Mostly S-phase | M-phase |
| Spatial restriction | None | Progenitors touching the ventricular wall (VZ) | Progenitors within electrical field (VZ + SVZ) | Progenitors touching the ventricular wall (VZ) |
| Label duration | Permanent if no further cell division | Promoter-dependent | Promoter-dependent | Max. 3 weeks |
| Dilution upon cell division | Yes | No | Yes (if non-integrating plasmid) | Yes |
| Detection of labeling | Immunohistochemistry and denaturation required | None required. Signal can be enhanced through immunohistochemistry | None required. Signal can be enhanced through immunohistochemistry | None required. Signal can be enhanced through immunohistochemistry |
| Sorting of live cells | No | Yes | Yes | Yes |
| Live imaging | No | Cells can be followed for long time (hours-days) | Cells can be followed for long time | Signal bleaches fast, spinning disk microscopy required |
| Delivery of genes | No | Possible | Possible | No |

**TABLE 2.** Troubleshooting table

| Step | Problem | Possible Reasons | Solution |
| --- | --- | --- | --- |
| 11, 13, 17 | Abortion | Maternal stress | Carefully handle analgesia after surgery<br>Provide additional analgesia when necessary<br>House animals in quiet area |
| | | Constriction of uterine horn during surgery | If uterine horns are not sufficiently relaxed, add 50 to 100 μl of 2 mg/ml magnesium sulphate on the uterine horns and rinse immediately with pre-warmed saline |
| | | Damage to uterine horn during surgery | Excessive pipette diameter. Keep the exterior diameter between 30 to 60 μm |
| 12 | Hydrocephaly | Too much FT injected | Reduce the volume of injection and/or the speed of injection. |
| 12 | Cell death (local necrosis) | DMSO impurity | Use DMSO provided in the CFSE kit or cell culture grade DMSO |
|  |  | Too much FT injected | Decrease injection volume |
| 12 | FT signal not visible in VZ progenitors of interest following labelling | Concentration of FT inadequate | Check FT concentration |
|  |  | Suboptimal site of injection (lateral vs third vs fourth ventricle) | Inject in ventricle closest to collection site |
|  |  | Injection outside of the ventricular system | During injection, make sure Fast Green is visualised in both lateral ventricles |
|  |  | Pipette tip blocked | Check pipette patency before injecting. If necessary, change pipette |
| 19 | Suture not intact during postoperative care | Knots were not tight enough or mouse removed the sutures | Re-suture under anaesthesia, or use surgical staples |

#### BOX 1 Mode of action of CarboxyFluorescein Diacetate Succinimidyl Esters for cell labeling

Carboxyfluorescein diacetate succinimidyl ester (CFDA-SE) is a commonly used reagent for long term fluorescent labeling of cells and flow cytometry-based proliferation studies^16,17^. It is a non-fluorescent molecule, which is highly membrane permeant due to its acetate groups. Upon cell entry, intracellular esterases cleave the acetate groups, resulting in a fluorescent and membrane non-permeant carboxy fluorescein succinimidyl ester (CFSE) that is trapped inside the cell. The succinimidyl chains of CFSE covalently bind to the amino groups of intracellular proteins, leading to long-lived labeling of the cell. At each cell division, CFSE is divided between the progeny, resulting in dilution of the signal^17^.

Once inside the cell, FT is diluted at each cell division, such that FT intensity can in principle be used to determine the number of cell divisions that have occurred between labeling and collection. This principle has long been applied in the hematopoietic system^15^, and can, to a certain extent, be applied in the CNS to distinguish neurons born directly from the VZ from those born indirectly via intermediate progenitors in the SVZ^7^. We established this principle in the neocortex using chronic IP BrdU perfusion from the time of FT injection to label all cells that entered S-phase at any time point after FT labeling (Fig 2a, b). When examined at P7, only 10% of the brightest FT^+^ cells were also BrdU^+^, indicating that the brightest FT^+^ cells are overwhelmingly born from VZ progenitors at the time of labeling (Fig 2c).

**Fig 2.**
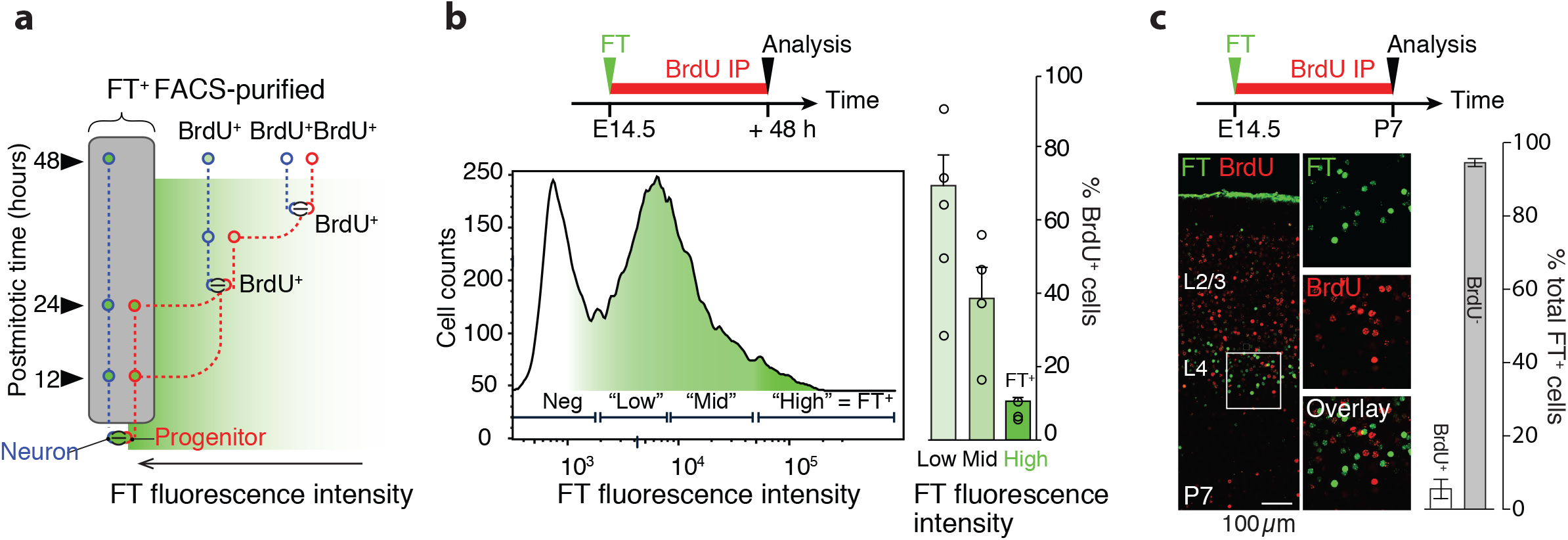
Brightly-labeled FT neurons (FT^+^) are directly born from VZ progenitors. **(a)** FT pulse-labeling combined with chronic BrdU perfusion allows labeling of cells that enter S-phase at any time after FT injection. Cells directly born from the VZ are identified as FT^high^ (“FT^+^ cells”,= brightest 10 % of cells) and BrdU^-^. **(b)** More than 90% of FT^+^ cells are BrdU^-^ when examined at 48 h after labeling by FACsorting, confirming that they were born directly from VZ progenitors at the time of FT labeling. **(c)** When examined at P7, the vast majority of FT^+^ cells are BrdU^-^, showing that FT^+^ cells are overwhelmingly born from VZ progenitors at the time of labeling. Illustrations from ref. 7.

To directly compare FT birthdating with BrdU birthdating, we pulse-injected BrdU at the time of FT labeling. While a broad set of S-phase progenitors in the cortical VZ and SVZ is labeled by BrdU, FT specifically labels M-phase cells at the ventricular border, such that these two markers label essentially mutually exclusive populations of cells (Fig 3a). When examined at P7, daughter FT^+^ neurons are found in a thin neocortical sub-layer, reflecting their identical time of birth, while daughter BrdU^+^ neurons have a broader laminar distribution, reflecting diverse cells of origins (VZ and SVZ) and developmental trajectories (Fig 3b). Similar results were obtained when comparing FT with *in utero* electroporation: 24 h after electroporation of a GFP-expressing plasmid in the neocortex, fluorescently-labeled cells were dispersed throughout the VZ, SVZ and intermediate zone (IZ), reflecting the broad and unsynchronized nature of targeted cells, as has been previously reported by electroporating BrdU-labeled plasmids^14^. In contrast, FT^+^ cells labeled at the same time were visible as a synchronized cohort, with strongly-labeled FT^+^ cells at the SVZ/IZ border and more weakly-labeled FT^+^ cells (most probably recycling progenitors) in the VZ (Fig 3c).

**Fig 3.**
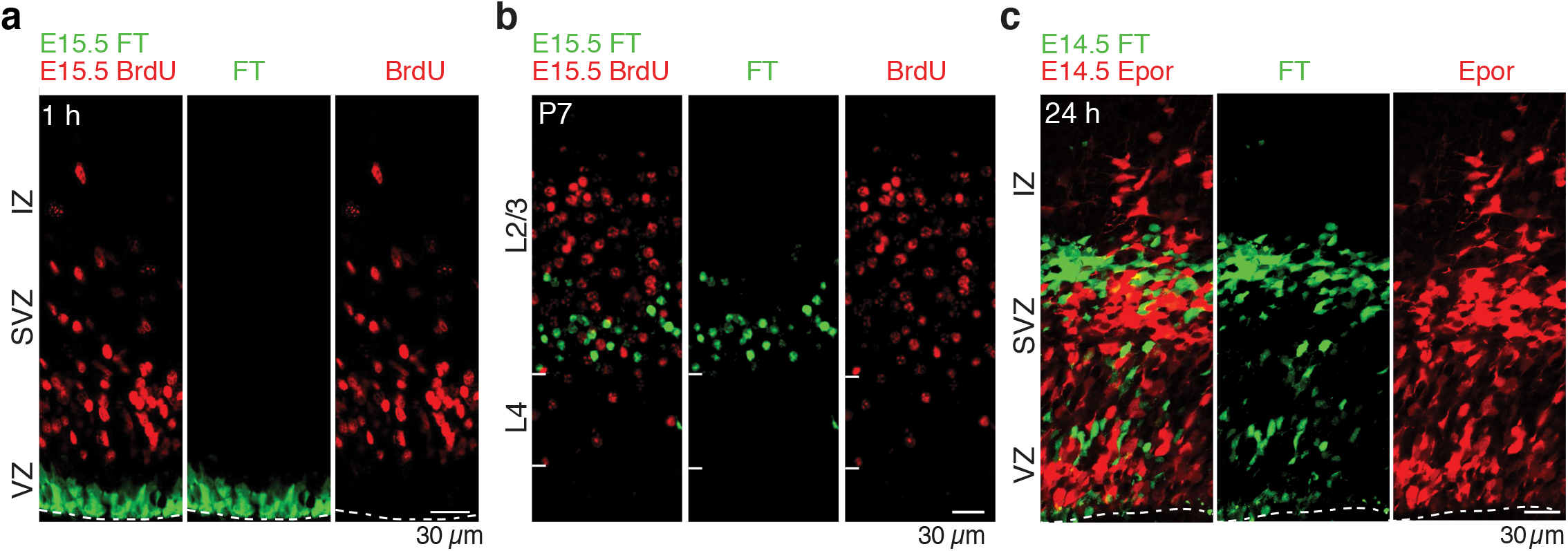
Comparison of BrdU, FT, and *in utero* electroporation pulse-labeling. **(a)** Simultaneous FT (intraventricular) and BrdU (i.p.) pulse-injection labels distinct cell populations. FT specifically labels mitotic progenitors in the VZ, while BrdU labels cells in S-phase across the VZ, SVZ and IZ. **(b)** At P7, FT^+^ cells form a thin sublayer in the neocortex, while BrdU^+^ cells are radially dispersed, reflecting their comparatively heterogeneous time and places of birth. **(c)** Simultaneous FT injection and in utero electroporation labels distinct cell populations. 24 h after electroporation with a reporter plasmid labeled cells are dispersed throughout the VZ, SVZ and IZ, while FT^+^ cells (*i.e*. daughter cells) are visible as a synchronized cohort at the SVZ/IZ border and more weakly-labeled FT^+^ cells (*i.e*. recycling progenitors) in the VZ. As in (b), the comparative heterogeneity of cells labeled by electroporation reflects heterogeneous place of birth, since electroporation affects progenitors across the VZ and SVZ (see *e.g.* Supplementary Fig. 1 in ref. 14).

### Potential further applications of FlashTag

Although within the CNS FT has been best characterized in the neocortex of the mouse, it can be applied to other brain regions and, in principle, can be used to birthdate and isolate any type of neuron that is born in contact with the ventricular space (or, potentially, any other cavity with fluid turn-over, such as the eye) throughout the brain and spinal cord, including in adult neurogenesis, across species, and in organoid preparations^18^. In addition, FT can also be used to label cells that migrate long distances from their place of birth, such as neurons of the olfactory bulb (Fig 4), and CFSE has been used to tag cells migrating through a given region by intraparenchymal injection^19^. The main limitations to this broad potential is the need to calibrate the half-life of FT in each condition, since temporal precision in tagging will depend on factors such as cerebrospinal flow rate, VZ permeability, and VZ progenitor kinetics. Finally, since fluorescence distributes into the cytoplasm, FT allows visualising the morphology of labeled cells, including, in some instances, projections (*e.g.* axons but also basal processes in radial glia).

**Fig 4.**
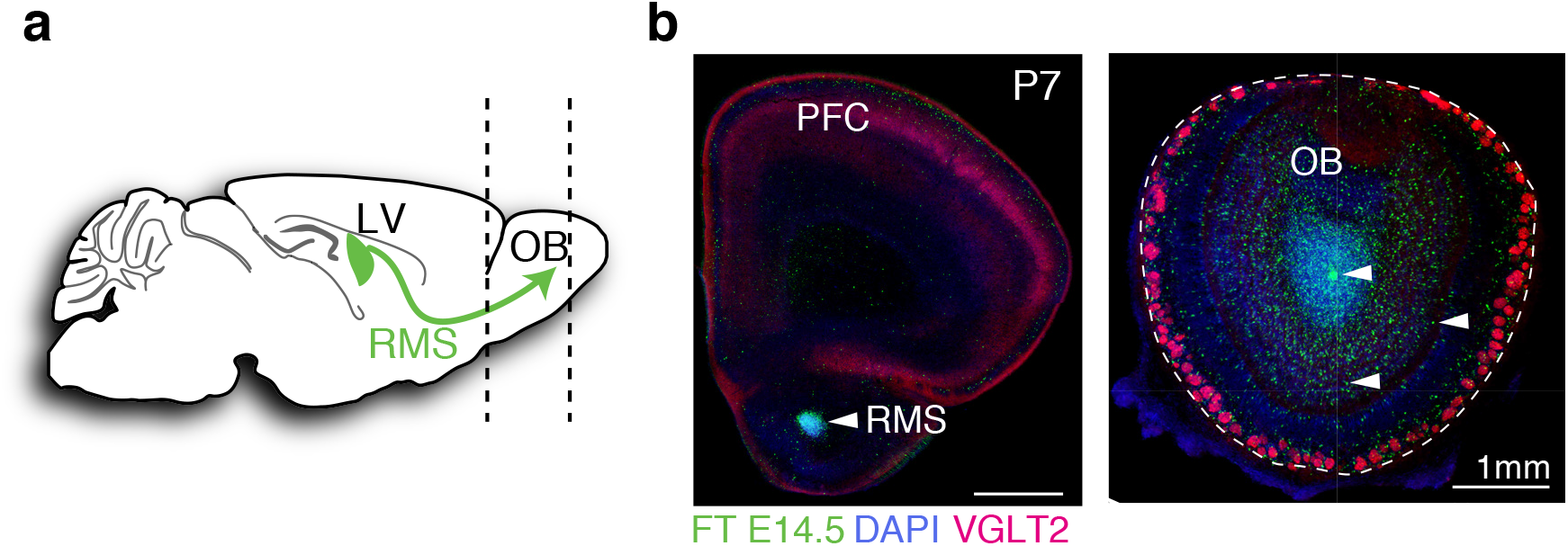
Labeling of remotely-migrating cells. **(a)** Neurons of the olfactory bulb are born at the border of the later ventricle and then migrate v*ia* the rostral migratory stream (RMS) to the olfactory bulb. **(b)** sections at the level indicated by the dashed lines in (a) reveals FT^+^ cells in the olfactory bulb (OB). Of note, cells in the pre-frontal cortex (PFC) do not migrate via the migratory stream.

Interestingly, by allowing the temporally precise labeling of sequentially-born neuronal populations, FT could be used as a template to establish correspondences between equivalent populations of cells across distantly-related species (*e.g.* mammals *vs.* sauropsids) without the need for potentially arbitrary molecular markers. By allowing isolation of isochronic neurons at precise sequential developmental stages, FT can also be used to directly characterise the transcriptional progression of cell populations over time and be used to validate or refine bioinformatics pseudotime alignment approaches^7,18^.

Finally, FT can be combined with other techniques, such as *in utero* electroporation, to specifically assess the effect of gene manipulation on a temporally restricted cohort of cells, as we have done in the past^7^. Of note, if combining FT with *in utero* electroporation, in the neocortex, on embryonic day 14.5, FT should be performed 4 – 6 h after electroporation to ensure double labeling. This interval corresponds to the time that electroporated cells need to reach the ventricular border (i.e. the S-to-M duration), and thus to be able to be labeled by FT. Importantly, the electroporation-to-FT delay should be calibrated for each structure and developmental stage to allow for optimal co-labeling.

### Limitations of FlashTag

- In the neocortex, FT signal is retained up until the third postnatal week. Over time, labeling becomes less bright, most probably due to protein turnover. However, when using CFSE, it is possible to enhance signal through immunohistochemistry using an anti-FITC antibody (see procedure and Fig. 6).
- FT signal intensity largely depends on the injection site: injection in the lateral ventricle will result in strong labeling in the forebrain but in a weaker signal in the mid- and hindbrain, while the converse is true for third ventricle injections. Thus, comparison of FT cell intensity across different brain regions (including across hemispheres) requires careful optimisation of injection site and volume, and calibration with chronic BrdU perfusion when focusing on direct neurogenesis from the VZ (see below). If comparing both cortices injection in the third ventricle is suggested to ensure equal labeling intensity in both hemispheres.
- When examining brains at early time points after FT labeling (*e.g.* 1 h), weak fluorescence can be observed in cells that were labeled by diffusion of CFSE into the VZ. A maximum intensity threshold can be applied to increase the signal to noise ratio in this case (Fig 6 A).
- Care should be taken when analysing cells close to the injection site, as contamination with unspecific labeling can occur along the needle tract.
- When examining labeled cells in the neocortex at P7, the top 10% brightest cells essentially correspond to cells that were directly born from the VZ at the time of FT labeling (*i.e.* only 10% of those are co-labeled by chronic BrdU perfusion, see Fig 2). As mentioned above, it is necessary to confirm this threshold in every examined brain structure and developmental stage by combining FT with chronic BrdU or EdU perfusion which, in our hands, is most reliably performed by using an intraperitoneally-implanted Alzet pump (see ref. ^7,20^). In addition, it should be noted that even when using chronic BrdU labeling, false positives are possible in the FT^+^ BrdU^+^ cells (*i.e.* these cells might wrongly be assumed to be born indirectly from the labeled progenitor yet be directly-born from a recycling progenitor at a later developmental stage).
- FT signal bleaches very fast, and thus the use of a spinning disk confocal is required when performing live imaging of FT labeled cells. Furthermore, it should be noted that when using FT to isolate cells for cell culture, the signal disappears substantially faster than *in vivo*.

## MATERIALS

### REAGENTS

CellTrace™ CFSE Cell Proliferation Kit (Thermo Fisher, cat. no. C34554) CytoTell™ Blue (AAT Bioquest, cat. no. 22252)

Note: We have validated the use of CellTrace™ CFSE and CytoTell™ Blue for FT labeling. Other compounds with the same working principle exist, but need to be validated before use: CellTrace™ Blue (Thermo Fisher, cat. no. C34574), CellTrace™ Violet (Thermo Fisher, cat. no. C34571), CellTrace™ Yellow (Thermo Fisher, cat. no. C34570), CellTrace™ Far Red (Thermo Fisher, cat. no. C34572); CytoTell™ Green (AAT Bioquest, cat. no. 22253), CytoTell™ Orange (AAT Bioquest, cat. no. 22257), CytoTell™ Far Red 650 (AAT Bioquest, cat. no. 22255), CytoTell™ UltraGreen (AAT Bioquest, cat. no. 22240). CytoTell™ Red 590 (AAT Bioquest, cat. no. 22261) did not in our hands work for FT labeling.

DMSO (Invitrogen, cat. no. D12345)

Fast Green (Sigma-Aldrich, cat. no. F7258)

Alzet osmotic pumps (Alzet, cat. no. 1003D)

PBS, 10x (Sigma-Aldrich, cat. no. D8662)

Saline, pre-warmed at 37°C

0.03 mg/ml Buprenorphinum

Chlorohexidine

Eye gel

Isoflurane

BrdU (Invitrogen, cat. no. B23151)

EdU (Invitrogen, cat. no. A10048)

Optional: 2 mg/ml Magnesium sulphate 1x HBSS (Gibco, cat. no. 14175-03)

TrypLE (gibco, cat. no. 12605-010)

Agarose Low Melt (Roth, cat. no. 6351.2)

Draq7 dye (Abcam, cat. no. ab109202)

Fetal Bovine Serum (Gibco, cat. no. 10082139)

Anti-Fluorescein antibody (abcam, cat. no. ab19491) Anti-rabbit fluorescent secondary antibody

Normal Horse Serum (Thermo Fisher, cat. no. 31874)

Hoechst (Thermo Fisher, cat. no. H3570)

Triton-X 100 (Sigma, cat. no. X100-100ML)

Fluoromount (Sigma, cat. no. F4680)

24 well plates

Glass slides for microscopy Painting brush for mounting

#### Mice

FT labeling has been performed in CD1 mice; other strains may be used, but adjustments in injection volume may be needed. **CAUTION** Experiments using live rodents must conform to all relevant governmental and institutional regulations.

### EQUIPMENT

Glass micropipette puller (Sutter Instrument, cat. no. P-97)

Glass capillaries (Drummond, cat. no. 3-000-203-G/X)

Surgery tools

Heating pad, 37 °C (Harvard Apparatus, cat. no. 340925)

Glass micropipette beveller (Sutter Instrument, cat. no. BV-10) Stereomicroscope (Leica, #M165 FC)

Dry bead sterilizer (Sigma, cat. no. Z378550)

NanoInject II auto nanoliter injector (Drummond Scientific, cat. no. 3000204)

Water bath

Cold light source (VWR, cat. no. 631-1085)

Optional : Cold light source with fiber optic light guide (Leica CLS100X)

Laboratory animal anesthesia system (VetEquip)

Centrifuge (VWR, cat. no. 521-1894)

Vibrating blade microtome (Leica, cat. no. 14047235613)

Dissection tools

Embryo spoon (Roth, cat. no. TL85.1)

5 ml syringes (Braun, cat. no. 8508607N)

1 ml insulin syringes (Terumo, cat. no. BS-N1N2713)

Adhesive tape (Mefix, cat. no. 310500)

Needled suture thread (Ethicon, cat. no. MPV 493)

Cotton pads

Cotton swabs

Thermometer

70 µM cell strainer (Greiner Bio-One, cat. no. 542070)

1.5 ml sterile Eppendorf tubes

Sterile pasteur transfer pipettes (Sarstedt, cat. no. 86.1171.001)

Petri dishes (Nunclon, cat. no. 150350)

### REAGENT SETUP

#### BrdU stock

Prepare 16 mg/ml BrdU in 50% DMSO and 50% H2O. Dissolve well and aliquot. Aliquots can be stored at −20°C for up to 1 month.

#### EdU stock

Prepare 10 mg/ml EdU in 50% DMSO and 50% H2O. Dissolve well and aliquot. Aliquots can be stored at −20°C for up to 1 month.

#### Fast Green

Dissolve Fast Green in 1× DMSO to a final concentration of 0.01% (wt/vol).

#### FlashTag working solution (CFSE)

Add 8 µl DMSO and 1 µl of Fast Green to one vial of Cell Trace™ CFSE for a final concentration of 10 mM. Working solution can be prepared in advance and stored at −20°C. **CRITICAL STEP** Use DMSO provided in the kit. Changing CFSE concentration can result in change in fluorescence levels of cell labeling and excessive concentration of CFSE can lead to cell death.

#### FlashTag working solution (Cytotell Blue)

Add 450 µl DMSO and 50 µl Fast Green to one vial of CytoTell™ Blue. Make 10 µl aliquots to avoid repeated freeze thaw cycles. **CRITICAL STEP** Use *in vitro* grade DMSO.

#### Optional: 2mg/ml Magnesium Sulphate

Dissolve 10mg of magnesium sulphate in 5ml of sterile water and sterilise using a filter.

#### Wash buffer

Prepare 0.01 % Triton-X 100 in 1X PBS.

#### Hoechst stock

prepare 10 mg / ml stock in 1X PBS.

#### Optional: Blocking-Permeabilisation buffer

Prepare 5 % Normal horse serum (NHS) with 0.3% Triton-X 100 in PBS.

### EQUIPMENT SET-UP

#### Alzet pump • TIMING 4 hours or overnight

Fill Alzet pump with 100 μl of 16 mg/ml BrdU or 10 mg/ml EdU with the help of a syringe and the provided blunt needle. Place pump in a sterile Eppendorf containing saline and incubate for at least 4 hours or overnight at 37°C to induce release. **CRITICAL STEP** If pump induction is skipped, release will be not stable over the first 4 hours. If chronic perfusion is required for more than 72 h, a new pump needs to be added 72 h after perfusion start to replace the previous one.

#### Pipette preparation • TIMING 5 min per pipette, usually done in batches ahead of time

Pull a glass capillary with the glass micropipette puller and cut the tip of each side to obtain an external diameter of 40 – 60 µm. Bevel each micropipette to an angle of 45-60 degree inclination. **CAUTION** Excessive pipette tapering can lead to pipette breaking and damage to uterine horn.

### PROCEDURE

#### Surgery preparation • TIMING 10 min

**1** Sterilize surgical instruments using a dry bead sterilizer.
**2** Pre-warm sterile saline at 37°C.

#### Surgical procedure • TIMING 20-30 min per mouse

**3** Place the mouse in the anaesthesia induction box and anesthetise using appropriate isoflurane and oxygen levels.
**4** Place the mouse on its back on a heated surgery platform with the snout inside the inhalation tube. **CRITICAL STEP:** The surgery should be finished within 20 - 30 minutes from this point.
**5** Apply eye gel to the eyes of the mouse to prevent them from drying out during the surgery.
**6** Fix the hind paws on the surgery heating pad using adhesive medical tape. **7|** Inject analgesic intraperitoneally (100 µl of 0.03 mg/ml buprenorphine). **8|** Disinfect the abdomen using a tissue soaked with Chlorohexidine.
**9** Make a midline incision of approximately 1-2 cm to expose the uterine horns. Place a sterile cotton pad on the lower part of the abdomen and moisturise with warm saline.
**10** With the help of a cotton swab carefully take the uterine horns out of the abdominal cavity and place them on the cotton pad. **CRITICAL STEP** i) Handle the uterine horns carefully and apply only minimal pressure. ii) Keep the uterine horns moist throughout the surgery using lukewarm saline.
**11** Optional: If the uterine horns are contracted they can be relaxed by applying 2-3 drops of 2 mg/ml Mg2SO4 solution. Instantly rinse the solution off with warm saline. **CRITICAL STEP** i) Use of Mg2SO4 is recommended only for embryos younger than E14.5. ii) If Mg2SO4 is not instantly washed away with saline it can cause damage to the uterine horns and result in abortions. iii) High concentration and volume of Mg2SO4 can result in abortions.
**12** Inject 0.5 µl of FT into the ventricle using a nanoinjector. Injection can be performed in the lateral or the third ventricle depending on the structure of interest (for strong labeling in the cortex embryos older than E14.5 should be injected in the lateral ventricle). **CRITICAL STEP** i) Excessive volume or speed of injection can result in hydrocephaly or death of embryo. ii) Pierce the uterine horn slowly and gently to avoid damage to the uterine wall as it can lead to abortion iii) Using mouth pipetting techniques will likely result in variable volumes of injection and comparison of FT intensities across embryos will thus be unreliable.
**13** Place the uterine horns carefully back into the abdominal cavity.
**14** Fill the abdominal cavity with lukewarm saline.
**15** Carefully place alzet pump into abdominal cavity for chronic BrdU or EdU administration.
**16** Close the peritoneal wall and then the skin using absorbable suturing thread. Alternatively, metallic clips can used.
**17** Inject analgesic i.p. (100 µl of 0.03 mg/ml temgesic) and return the mouse to its cage.
**18** Place the cage on a heating pad. The mouse should recover within a few minutes.

### Postoperative care

**19** Check the mouse during the next two days for any signs of discomfort and check if the sutures are intact. Depending on local guidelines, additional analgesia can be administered if signs of discomfort are present.

### FACsorting FT^+^ population from FT-injected embryos• TIMING 120 min

**20** Fill 4 petri dishes with HBSS and place them on ice.
**21** Sterilize all dissecting tools with 70% ethanol.
**22** Prepare 4% low-melt agarose in HBSS.
**23** Set the vibrating microtome temperature to 4 °C or cool the cutting chamber with ice and fill with ice cold HBSS.
**24** Sacrifice the dam by cervical dislocation following deep anaesthesia and make a midline incision in the abdomen to access the embryos. With the help of fine forceps pinch open the uterine horns and collect the embryo heads into a petri dish filled with HBSS placed on ice.
**25** Melt the 4% agarose and let cool down to 42 °C while proceeding to the next step.
**26** Screen the heads for FT signal using appropriate fluorescence filters and dissect FT^+^ brains (Fig 5a, b). **CRITICAL STEP** Keep the brains in HBSS on ice whenever possible to minimise cell death.

**Fig 5.**
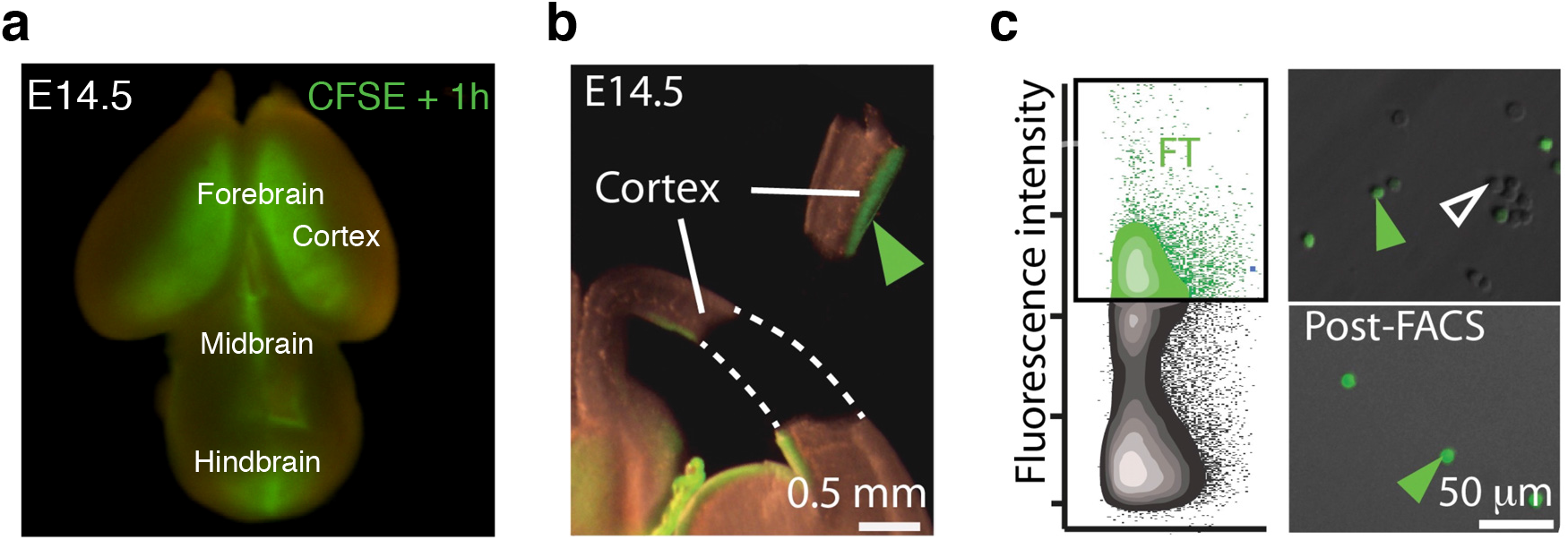
Detecting FT labeling in live tissue. **(a)** 1 hour after FT labeling at E14.5, strong fluorescence is seen in whole-mount preparations, delineating the ventricular system. **(b)** Upon microdissection, strong fluorescence is visible at the apical border of the tissue sample. **(c)** FAC-sorting of FT^+^ cells. Only cells with high levels of fluorescence intensity are sorted (closed arrowheads), and FT^-^ cells are excluded (open arrowhead). Illustrations in b and c from ref. 7.

**27** Embed the brains in 42°C agarose using plastic moulds placed on ice.
**28** Once the agarose solidifies, carefully remove agar block from plastic mould. Using a blade, remove excess agarose and cut small blocks leaving no more than 3 mm of agarose around the brain.
**29** Glue the agarose blocks containing the tissue to the vibrating microtome support and cut 300 - 600 µM thick coronal sections with low speed and vibration. Collect slices in a petri dish with ice-cold HBSS using an embryo spoon.
**30** With the help of a stereomicroscope, dissect the regions of interest using micro-knives and transfer the dissected part to a 1.5 ml micro-centrifuge tube placed on ice with a plastic pipette. **CRITICAL STEP** Keep the tissue in HBSS on ice whenever possible to minimise cell death.
**31** Remove HBSS from the tube containing tissue pieces, leaving not more than 100 µl, and add 400 µl of 1x TrypLE. Incubate for 3 - 5 min at 37 °C in a water bath.
**32** Add 200 µl of FBS to inactivate the TrypLE. Centrifuge at 300 g for 5 min at room temperature, remove supernatant, add 500 µl of medium, triturate the tissue manually to make a homogenous single-cell suspension.
**33** Filter cells through a 70 µm filter into a sterile Eppendorf tube and add Draq7 to identify dead cells and place Eppendorf tube on ice.
**34** Sort cells with a Beckman Coulter Moflo Astrios FAC-sorter at 4°C. Singlet cells can be defined using the forward and side scattering properties (FSC and SSC). Eliminate dead cells using Draq7TM signal. CFSE fluorescence emission can be measured around 530 nm (Band Pass Filter 530/30). Collect the top 5^th^-10^th^ percentile of cells based on the SSC-A/CFSE density plot (Fig. 5c).

### Immunohistochemistry of FT labeled brains • TIMING 3 days

#### Day 1: Dissection and fixation of brains

**35** For embryonic tissue: Dissect brains in ice-cold HBSS solution and fix in 4 % PFA overnight. Remove PFA and replace with 1x PBS. It is also possible to first fix the entire head before dissecting the brain.
**36** For postnatal tissue: Perfuse deeply anesthetised mouse with 0.9 % NaCl, followed by 4 % PFA. Dissect brain and postfix in 4% PFA overnight. Remove PFA and replace with 1x PBS.

#### Day 2: Sectioning with vibrating microtome

**37** Embed fixed brains in 4 % agarose. Cut small blocks leaving around 3 mm of agarose on each side of the brain.
**38** Fill the vibrating microtome with 1x PBS (optional: use ice-cold PBS), set the blade at 90 degrees, and cut 60 µm thick sections. Collect sections in 24 well plates filled with 1x PBS. Note: Sections can be stored for several days at 4°C. The FT signal is stable for several days, but the integrity of tissue can be compromised when stored for a long time.
**39** Optional: Incubate sections in blocking-permeabilization buffer for 40 min at RT.
**40** Prepare primary antibody: dilute anti-Fluorescein antibody 1:1000 in antibody dilution buffer. Incubate sections with primary antibody overnight on a shaker at 4°C.

#### Day 3: Immunohistochemistry against CFSE to enhance signal (Fig. 6a)

**41** Remove antibody solution and wash sections 3x with 0.1% PBST on a shaker.
**42** Prepare secondary antibody: dilute secondary antibody in dilution buffer. Incubate sections with secondary antibody for 1-2 hours on a shaker at RT.
**43** Optional: Label nuclei with Hoechst. Dilute Hoechst 1:5000 in 1x PBS. Incubate the sections with Hoechst for 5 min at RT on a shaker.
**44** Wash sections 3x with washing buffer on a shaker (5 min at RT each).
**45** Mount sections onto a glass slide using a brush. Let the sections dry and then coverslip with Fluoromount.

**Fig 6.**
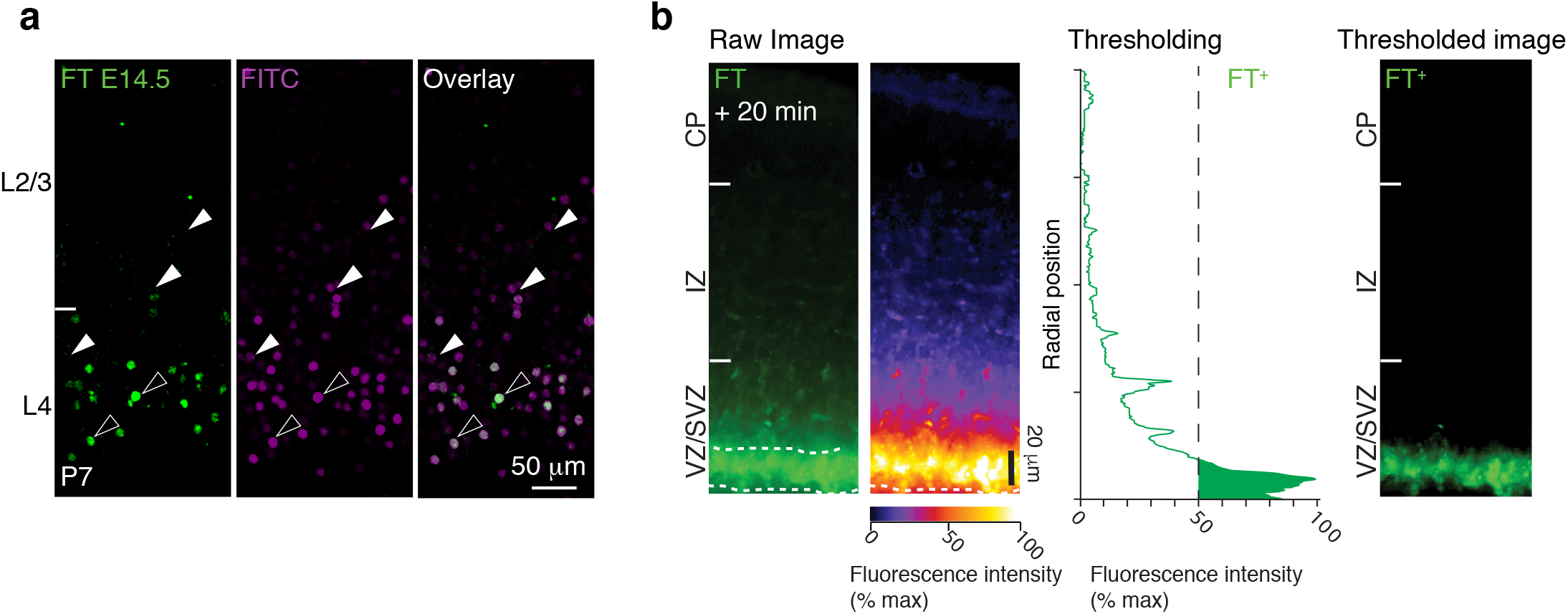
Detecting FT labeling in fixed tissue. (**a**) Anti-FITC antibody can be used to increase CFSE signal in the neocortex. Open arrowheads show high FT^+^ cells co-labeled with anti-FITC, and closed arrowheads show cells that are not visible without antibody staining. (**b**) Twenty minutes after FT labeling strong fluorescence is observed in cells in contact with the ventricular wall, while weaker fluorescence (likely due to diffusion of CFSE into the tissue) is observed in the adjacent cell layers of the VZ. A maximum intensity threshold is applied to reveal high FT^+^ cells (50 % of maximum signal). (**b**) Anti-FITC antibody can be used to increase CFSE signal in the neocortex. Open arrowheads show high FT^+^ cells co-labeled with anti-FITC, and closed arrowheads show cells that are not visible without antibody staining.

#### Day 4: Imaging and thresholding for high FT^+^ population (Fig. 6b)

**46** Acquire images with confocal microscopy using the appropriate excitation:

| Labeling | Excitation/Emission |
| --- | --- |
| Cytotell Blue | 403/454 |
| CellTrace CFSE | 495/519 |

**47** When acquiring the images maintain the entire dynamic range (allowing only few over- and under-saturated pixels).
**48** Adjust contrast during image post processing to highlight the top 50% brightest to identify the high FT^+^ population (i.e, the cells that were labeled by mitotic uptake of FT and that did not undergo subsequent cell division).

### TIMING

Steps 1-2, surgery preparation: 10 min

Steps 3-18, surgical procedure: 20 – 30 min per mouse

Step 19, postsurgery animal monitoring: 48 h

Steps 20 – 34, FACs-sorting FT^+^ cells, 120 min

Steps 35 – 36, dissection and fixation of brains, 1 day

Steps 37 – 40, vibratome sectioning and incubation with primary antibody, 1 day

Steps 41 – 48, incubation with secondary antibody and imaging, 1 day

## ANTICIPATED RESULTS

Cells born in the ventricular zone across the CNS can be easily followed from birth until at least 3 weeks postnatally. Examples of typical FT labeling are presented in Figure 3 (neocortex) and Figure 4 (olfactory bulb). Labeled cells can be isolated with FACS for various applications, including single-cell RNA sequencing, or live tissue can be collected for live imaging or electrophysiology. A maximum intensity threshold can be used to strongly enrich in cells that were directly born from the VZ at the time of FT labeling, both when isolating cells using FACS, as well as when analysing labeled cells with histology.

## Acknowledgements

We thank the members of our laboratory for helpful discussions. We also thank A. Benoit and M. Lanzillo for technical assistance, L. Telley for his contribution to initial discussions and R. Wagener for providing the photomicrograph in Fig. 4. Work in the Jabaudon laboratory is supported by the Swiss National Science Foundation, the Fondation des Hôpitaux Universitaires de Genève and the Brain and Behavior Foundation.

## Author Contributions

S.G. and D.J. developed the initial protocol which was later updated by members of the laboratory including P.O., S.G. wrote the manuscript with the help of P.O. and D.J., S.G. performed the experiments.

